# Phellem translational landscape throughout secondary development in *Arabidopsis* roots

**DOI:** 10.1101/2021.02.08.429142

**Authors:** Ana Rita Leal, Pedro Miguel Barros, Boris Parizot, Helena Sapeta, Nick Vangheluwe, Tonni Grube Andersen, Tom Beeckman, M. Margarida Oliveira

## Abstract

The phellem is a specialized boundary tissue providing the first line of defense against abiotic and biotic stresses in organs undergoing secondary growth. Phellem cells undergo several differentiation steps, which include cell wall suberization, cell expansion and programmed cell death. Yet, the molecular players acting particularly in phellem cell differentiation remain poorly described, particularly in the widely used model plant *Arabidopsis thaliana*.
Using specific marker lines we followed the onset and progression of phellem differentiation in *A. thaliana* roots, and further targeted the translatome of new developed phellem cells using Translating Ribosome Affinity Purification followed by mRNA sequencing (TRAP-SEQ).
We showed that phellem suberization is initiated early after phellogen (cork cambium) division. The specific translational landscape was organized in three main domains related to energy production, synthesis and transport of cell wall components, and response to stimulus. Novel players in phellem differentiation, related to suberin monomer transport and assembly, as well as novel transcription regulators were identified.
This strategy provided an unprecedented resolution of the transcriptome of developing phellem cells, giving a detailed and specific view on the molecular mechanisms controlling cell differentiation in periderm tissues of the model plant *Arabidopsis*.

**Significance statement:** To improve the understanding of phellem differentiation into a suberized protective layer, we followed the establishment of periderm in *Arabidopsis* roots and sequenced the phellem-specific translatome. We found that phellem suberization occurs shortly after pericycle cell divisions with the induction of pivotal suberin biosynthesis genes. In parallel, we detected the activation of three central genetic modules acting throughout the phellem differentiation. This study provides a unique and targeted genetic resource for further functional studies of phellem tissues.

## Introduction

Adaptation and evolution of plants to the terrestrial environment was made possible by the development of specialized lipid and phenolic barriers that improved resistance to dehydration and protection against other environmental threats. In mature stems and roots undergoing secondary development, as well as in tubers and fruits, this protection is mostly provided by a suberized periderm (Ragni and Greb, 2018; Evert, 2006). The periderm is a complex structure, organized in three specialized layers: phellogen, phelloderm, and phellem. The phellogen (or cork cambium), is the meristematic layer that dedifferentiates from parenchymatous cells during secondary development. Periderm development occurs through phellogen activity, producing phelloderm cells towards the inside, and phellem cells towards the outside (Evert, 2006; Pereira, 2007). While phelloderm cells differentiate into parenchyma, phellem cells undergo several differentiation steps, which include cell wall suberization, cell expansion and programmed cell death (Evert, 2006; Crang *et al.,* 2018). As the outermost layer, phellem provides the ultimate protection in diverse plant organs.

The molecular regulation of periderm development has been studied in several plant species through transcriptomic analyses (Vulavala *et al.,* 2017; Soler *et al.,* 2011; Ginzberg *et al.,* 2009; Soler *et al.,* 2007; Rains *et al.,* 2018; Boher *et al.,* 2018; Alonso-Serra *et al.,* 2019; Lopes *et al.,* 2019). Collectively, these studies highlighted multiple genes related to phellem differentiation and more particularly to the suberization process, which is shown to be highly conserved across different species. Suberin is a complex lipophilic macromolecule composed of a diversity of aliphatic and aromatic components (monomers) that are covalently linked by primarily ester bounds (Bernards, 2002; Graça, 2015). Core pathways required for suberization retrieved in those transcriptomic studies include genes involved in the synthesis of specific aliphatic *(Long-Chain acyl-CoA Synthetase 2 - LACS2, Fatty acyl-coenzyme A reductase 1/4/5 - FAR1/4/5*, Cytochrome P450 86A1 - *CYP86A1* and *CYP86B1*) and aromatic components *(phenylalanine ammonia lyase - PAL, Cinnamate-4-hydroxylase - C4H* and *Ferulate 5-hydroxylase - F5H*), assembly (*Glycerol-3-phosphate Acyl-Transferase 5/7* - *GPAT5/7*, *Aliphatic suberin feruloyl transferase* - *ASFT* and Fatty alchol:caffeoyl-CoA transferase- *FACT*) and transport *(ATP-binding cassette G2/6/20* - *ABCG2/6/20*) of monomers (Vishwanath *et al.,* 2015; Graça, 2015; Philippe *et al.,* 2020). Previous transcriptomic studies on cork oak (Teixeira *et al.,* 2018) and potato (Vulavala *et al.,* 2019) using micro-dissected periderm samples (including phellogen and early-developed phellem) detected expression of suberization-related genes, suggesting that this process occurs at an early stage of periderm development. Expression activity of multiple transcription factors (TFs), including members of the MYB, NAC and WOX families, was also detected across targeted phellem transcriptomes, suggesting a conserved regulatory landscape acting during phellem development. Overall, these genome-wide transcriptomic studies greatly improved the understanding of the genetic mechanisms contributing to phellem differentiation.

Recently, the model plant *Arabidopsis thaliana* was proposed as a suitable model to study periderm development and differentiation (Wunderling *et al.,* 2018; Campilho *et al.,* 2020). As in woody stems, secondary development in *Arabidopsis* roots is organized in two concentric meristems: the inner vascular cambium and the outer phellogen. Morphological analysis of root development (Wunderling *et al.,* 2018), together with lineage tracing studies (Smetana *et al.,* 2019), confirmed periderm ontogeny from pericycle cells, that divide to give origin to the root phellogen. Initial pericycle cell divisions were reported already at 6 days after germination (DAG) in primary roots below hypocotyl-root junction, resulting in phellogen establishment (Smetana *et al.,* 2019). Core genes related to suberization are also strongly expressed in emerging phellem cells from *Arabidopsis* roots. Moreover, putative transcriptional regulators of phellem detected in woody species are also found in *Arabidopsis* phellem (Wunderling *et al.,* 2018), demonstrating once more the conservation of the developmental process of phellem between diverse species. In the present study we aimed to obtain a detailed view on the translatome of *Arabidopsis* phellem at initial stages of differentiation. After establishing a timeline for the onset of suberization in the root periderm, we performed Translating Ribosome Affinity Purification followed by mRNA sequencing (TRAP-SEQ) of newly formed phellem cells. TRAP-SEQ enabled for the first time the isolation of an mRNA pool from young phellem cells, avoiding the manual dissection of the tissue. The specific translational landscape of phellem cells revealed, not only previously described phellem molecular profiles, but also novel candidates related to synthesis and transport of suberin, as well as hormonal and transcriptional components putatively involved in phellem development. This strategy provided an unprecedented resolution of the transcriptome of developing phellem cells, giving a detailed and specific view on the molecular mechanisms controlling cell differentiation in periderm tissues of the model plant *Arabidopsis*. This study represents a starting point for in-depth analysis of the molecular regulation of phellem differentiation.

## Results

### Suberization timeline in phellem differentiation of the Arabidopsis root

To follow the establishment of new phellem cells, we performed a detailed morphological and spatio-temporal characterization of suberization during secondary development of *Arabidopsis* roots. We used previously established *Arabidopsis* suberin gene marker lines together with histological suberin staining to follow up phellem differentiation in transverse and longitudinal planes of roots. Promoter reporter lines for two suberin biosynthesis genes - *FAR4* (Domergue *et al.,* 2010) and *GPAT5* (Barberon *et al.,* 2016; Beisson *et al.,* 2007) - were used to monitor activation of suberin biosynthesis, while Nile Red and Fluorol Yellow (FY) were used for suberin detection in cell walls. FY (Brundrett *et al.,* 1991) and Nile Red (Ursache *et al.,* 2018) are highly sensitive fluorescent stains that can specifically label suberin-enriched cell walls in fresh or ClearSee treated roots, respectively. Observations were always conducted below root-hypocotyl junction, where secondary development is initiated (Smetana *et al.,* 2019), on time-course experiments starting 6 DAG to 11 DAG. At the earliest time-point, cross sections revealed cell divisions at procambium and at pericycle level (xylem and phloem pole) (Fig. **1a** – 6 DAG, arrows), confirming the onset of secondary growth and more particularly phellogen development. However, on these sections *FAR4* promoter activity was not detected in pericycle daughter cells nor in endodermal cells. Longitudinal imaging of the same root zone with the *pGPAT5::BLRP-FLAG-GFP-RPL18* marker line stained with Nile Red, revealed that suberin is present only in the endodermis cell walls (Fig. **1b**, S**1a**). As observed for *FAR4, GPAT5* expression was also not detected at this developmental stage (Fig. **1b**, S**1b**). Together, these observations indicate that the endodermis is not actively producing suberin at this stage, despite being already suberized, and the suberin biosynthesis markers are not active in this region of the root in any tissues. At 7 DAG we observed an increase in cell proliferation in the stele, including both in the procambium and the pericycle, likely imposing pressure on the overlying endodermis layer (Fig. **1a** – 7 DAG arrow), while cortex and epidermis maintained their normal cell shape. Punctual expression of *FAR4* was also detected in some cortex cells (Fig. **1a** – 7 DAG), in agreement with stochastic suberin staining detected in longitudinal imaging (Fig. **1b** – 7 DAG). At this stage suberin was no longer detected in the endodermis using Nile Red staining, which, however, was possible using FY staining in wild type roots (Fig. **S1a** – 7 DAG). At this time point, *GPAT5* was detected in the stele region of root, however, cell specificity could not be determined due to the low staining penetration across cell walls (Fig. **1b**, **S1b**). At 8 DAG, *FAR4* expression was detected specifically in pericycle-derived cells (Fig. **1a**). At the same time point, the *GPAT5* was detected in more cells and some of which were suberized, as revealed by Nile Red staining (Fig. **1b**). The appearance of these suberizing cells, with marker activation and suberin being detected, indicates that phellogen is active and phellem cells are being produced. The early activation of *GPAT5* promoter at 7 DAG, prior to suberin detection, suggests that this gene is a suitable marker for early stages of phellem development. Further, at 9 DAG, the number of differentiated phellem cells increased, as expression of both markers and suberization were detected in more cells (Fig. **1a, 1b**, **S1a**, **S1b**). After the initiation of phellem suberization, expansion of phellem cells was detected, together with abscission of cortex and epidermis layers, while the endodermis cells were no longer observed (Fig. **1a** **- 9DAG, S1c**). The number of circumference cells limiting the stele region, divided from pericycle cells (Fig. **1a** – 6DAG arrows), increased only till 7 DAG and this cell number was maintained in further analysed days (Fig. **S1c**), after detection of phellem as the limiting tissue. This demonstrates that phellem cell expansion starts in parallel with suberization. Finally, at 10 DAG, while *FAR4* and *GPAT5* expression was reduced, a continuous layer of suberized phellem cells could be identified (Fig. **1**, **S1a**, **S1b**). At this later stage, phellem is displayed as a continuous layer of suberized cells surrounding the root stele (Fig. **1a, 1c**) and in the longitudinal view, suberized phellem appears as a tissue covering the external part of the root, with cells organized in a compact and regular arrangement (Fig. **1 b**; **S1a**, **S1b**).

**Fig. 1.**
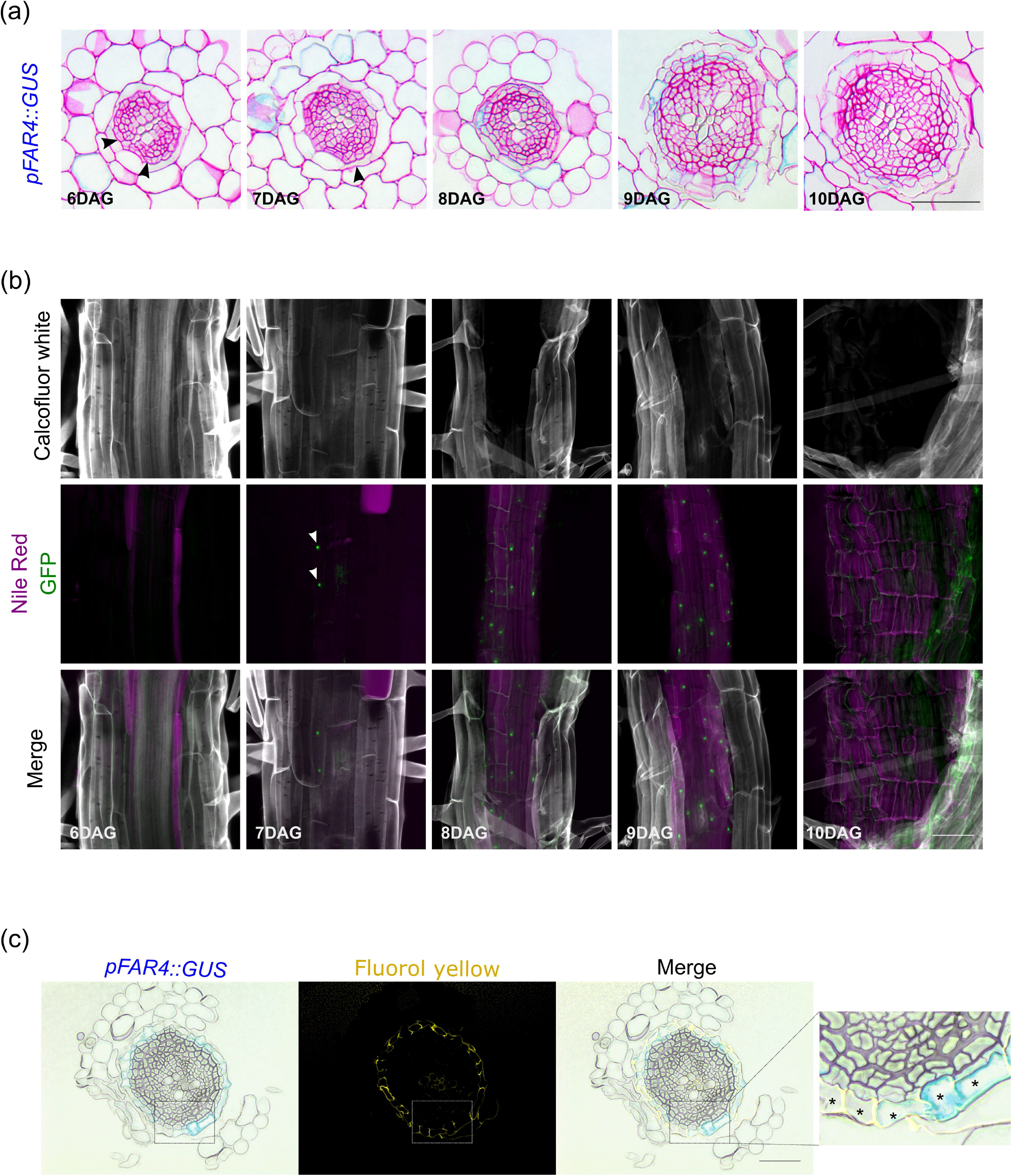
Activation of specific suberin biosynthesis markers during periderm development of Arabidopsis thaliana roots. (a) Plastic embedded cross-sections of FAR4::GUS taken below the hypocotyl-root junction, between 6 and 10 DAG. Arrowheads indicate new cell divisions in pericycle cells (6 DAG) and endodermis deformation (7 DAG). (b) GPAT5::BLRP-FLAG-GFP-RPL18 expression (green) in the phellem differentiation zone, below the root-hypocotyl junction, between 6 and 10 DAG. Cell walls (grey) visualized by calcofluor white and suberin (purple) using Nile red staining, after ClearSee treatment. Arrowheads indicate GPAT5::BLRP-FLAG-GFP-RPL18 expression. The 3D maximum projection figures were obtained by confocal laser scanning with Z-stack images. (c) Plastic embedded cross-sections of 11 DAG plants of FAR4::GUS taken below hypocotyl-root junction and stained with Fluorol Yellow for suberin. Asterisks (*) show suberized phellem cells. Scale bars: 50 μm. Images are representative of all analysed samples.

Collectively, this chronological study framed the onset of root phellem development between 7 to 8 DAG, just below the root-hypocotyl junction, closely followed by cell wall suberization of these newly formed cells.

### Isolation of phellem translatome

To identify the pathways mediating the early stages of phellem differentiation we performed a transcriptomic analysis at the onset of phellem suberization, which occurs, based on our microscopic study, at 8 DAG in the mature region of the root. We used the *pGPAT5::BLRP-FLAG-GFP-RPL18* line, where *pGPAT5* driven expression of the tagged Ribosomal Protein L18 *(RPL18)* allowed immunopurification and sequencing of the phellem-specific translatome (TRAP-SEQ). As a control, we used plants expressing the same construct under control of a near-constitutive promoter *(pUBQ10::BLRP-FLAG-GFP-RPL18)*, to obtain a representative transcriptome of all root cell types of the same root region and developmental stage (Fig. **2a**). Using Illumina sequencing, we obtained a total of 166 million reads from triplicate samples of the total root translatome (*UBQ10*-specific) and the suberizing phellem translatome (*GPAT5*-specific). A total of 5,385 genes were differentially expressed (adjusted p-value ≤ 0.01, |FC| ≥ 2) between *GPAT5*-specific and *UBQ10*-specific translatomes. From the detected differentially expressed genes (DEGs), 2,750 (51%) were more abundant in suberizing phellem (Table **S2**).

**Fig. 2.**
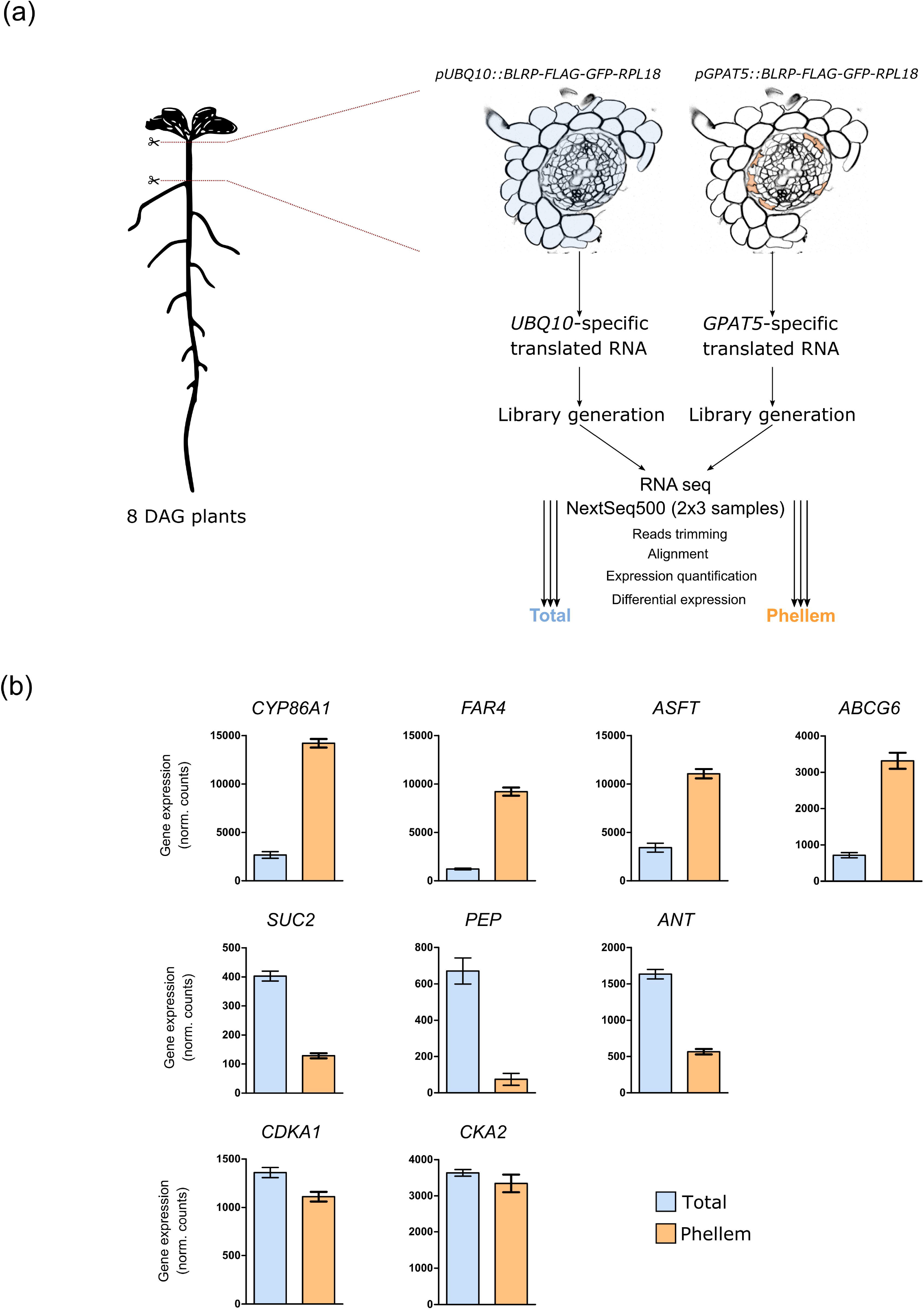
Isolation of specific Arabidopsis thaliana phellem transcripts. (a) Arabidopsis roots were collected from phellem development zone, from hypocotyl-root junction to the first emerged lateral root. The GPAT5 and UBQ10 promoters were used to drive expression of BLRP-FLAG-GFP-RPL18 cassette in phellem cells and in the total root cells, respectively. Immunocapture of ribosomes containing the fusion tag allowed cell-specific purification of mRNAs attached to these ribosomes followed by RNA sequencing. Biological triplicates were run for each condition. Reads from sequencing data were treated and differential expression of genes between whole root and phellem cells was calculated. (b) Transcript abundance of marker genes from collected phellem cells (GPAT5-specific) and total-root (UBQ10-specific): CYP86A1, FAR4, ASFT and ABCG6 are markers for cell-wall suberization; SUC2, PEP and ANT are markers for epidermis and cortex cells; CDKA1 and CKA2 are described as housekeeping genes. Data represent mean normalized counts ± standard deviation (n=3).

To assess the purity of the pull-down, we confirmed the expression profile of genes with known cell-specific localization in the *Arabidopsis* root (Fig. **2b**). Accordingly, genes involved in suberin monomer biosynthesis, *CYP86A1* (Höfer *et al.,* 2008)*, FAR4* (Domergue *et al.,* 2010), esterification, *ASFT* (Gou *et al.,* 2009), and transport *ABCG6* (Yadav *et al.,* 2014) were significantly enriched in the *GPAT5*-specific translatome. Moreover, genes specifically expressed in phloem *(SUC2)* (Truernit and Sauer, 1995), root cortex (*PEP*) (Mustroph *et al.,* 2009) and vascular cambium (*ANT*) (Randall *et al.,* 2015) were enriched in *UBQ10-specific* samples, and the expression of genes associated with housekeeping functions *CDKA;1* and *CKA2* remained unchanged in both translatomes. Also, the expression pattern of these selected genes assessed by RT-qPCR agreed with the RNA-seq data (Fig. **S2**), with both Log2FC showing a linear correlation coefficient of R^2^ = 0.96 (Fig. **S2**). To further validate the specific enrichment in phellem cells we studied the expression pattern of a GUS-reporter line for Nuclear Factor YC12 *(NF-YC12* - AT5G38140), which was detected to be enriched in the *GPAT5*-specific translatome (Table **S2**). In this line GUS expression was detected in secondary developed root zone 7 DAG, specifically in cells matching the localization of newly developed phellem (Fig. **S3**).

Overall, these results support the potential of the generated dataset in discriminating the suberizing phellem translatome from the global root translatome, at the onset of phellem development.

### Functional categorization of the phellem translatome

Functional enrichment analysis of the DEGs identified contrasting molecular signatures for *GPAT5*-specific and *UBQ10*-specific translatomes (Fig. **3**, **S4**; Tables **S3**). Regarding the phellem-specific translatome, we found a significant enrichment (FDR ≤ 0.05) of 277 GO terms, distributed among Cellular Component (13), Biological Process (117) and Molecular Function (147) categories (Fig. **3**; Table **S3A**). *Membrane* (GO:0016020) was highly overrepresented with 433 DEGs, followed by *Endomembrane system* (GO:0012505) and *Extracellular region part* (GO:0044421) with a lower number of associated DEGs. In the Biological Process category, *Response to stimulus* (GO:050896) was significantly enriched, with 404 DEGs. These were distributed among related terms, including *Response to abiotic stimulus* (GO:0009628), *Response to biotic stimulus* (GO:0009607), and *Response to hormone stimulus* (GO:0009725). More particularly, response to hormones included *Response to abscisic acid* (GO:0009737), *salicylic acid* (GO:0009751) and *jasmonic acid* (GO:009753) stimulus (Fig. **3**, Table **S3A**). *Developmental process* (GO:0032502), *Transport* (GO:0006810) and *Establishment of localization* (GO:0051234) terms are also represented, with each term holding approximately 10% of the DEGs annotated to Biological process terms. GO terms that relate to secondary metabolism were also enriched, namely *Phenylpropanoid metabolic process* (GO:0009698), *Suberin biosynthetic process* (GO:0010345) and *Cutin biosynthetic process* (GO:0010143). In the Molecular Function category, *Hydrolase* (GO:0016787), *Transferase* (GO:0016740) and *Oxidoreductase activity* (GO:0016491) terms were enriched, individually representing more than 10% of annotated DEGs. Lipid-related Molecular Function terms were also detected, including *Fatty acid synthase activity* (GO:0004312)*, Long-chain fatty acid-CoA ligase activity* (GO:0004467) and *Very long-chain fatty acid-CoA ligase activity* (GO:031957).

**Fig. 3.**
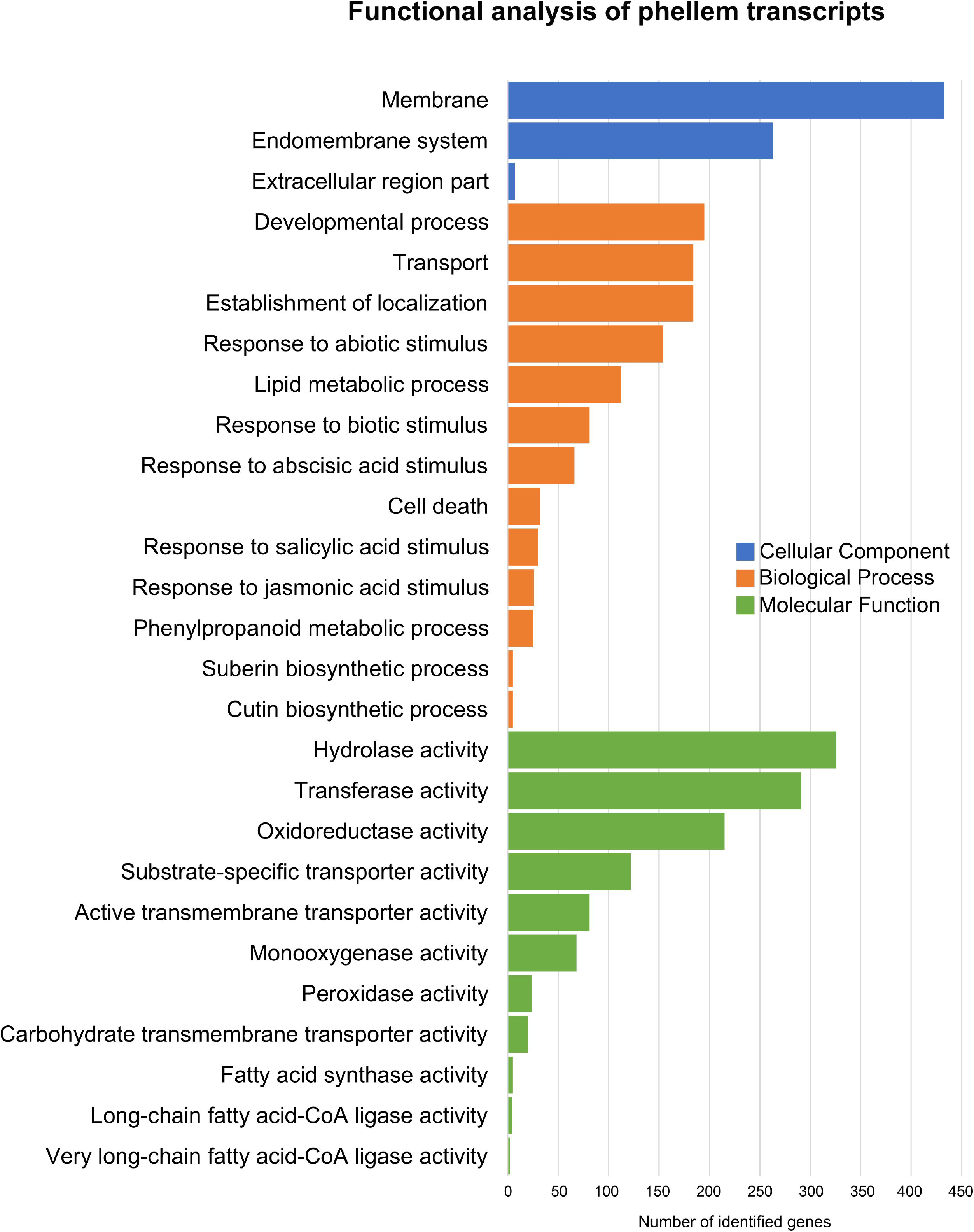
Representative GO terms significantly enriched within genes differentially expressed in phellem samples. Colored bars represent the number of genes in the selected GO terms in Cellular component, Biological process and Molecular function.

The *UBQ10*-specific translatome displayed a significant enrichment of 221 terms mostly distinct from the *GPAT5*-specific translatome. We found 8 functional categories were shared with the *GPAT5*-specific translatome, including (Fig. **S4**; Table **S3B**): Cytoplasm (GO: 005737), *Membrane* (GO:0016020), *Response to abiotic stimulus* (GO:0009628), *Response to water* (GO:0009415) and *Response to water deprivation* (GO:009414). Among other terms, the *UBQ10*-specific translatome showed a functional enrichment of: *Intracellular* (GO:0005622), *Organelle* (GO:0043226) and Protein complex (GO:0043234) from Cellular Component category; *Response to auxin stimulus* (GO:0009733), *Regulation of cell cycle* (GO:0051726) and *Regulation of meristem structural organization* (GO:0009934) from Biological Process; and *DNA binding* (GO:0003677) and *Structural molecule activity* (GO:0005198) for Molecular function category.

### Active metabolic pathways in phellem differentiation

Phellem-specific translatome revealed to be significantly distinct from the total root translatome, with a high number of DEGs enriched in this tissue. This is in line with previous analyses which revealed that phellem is the most transcriptional diverse tissue among a variety of other tissues, with the suberization processes being the most distinct characteristic of differentiating phellem (Lopes *et al.,* 2019; Alonso-Serra *et al.,* 2019). Considering this, we selected a short list of genes highly represented and with putative relevance for phellem differentiation in *Arabidopsis*. The selected genes are associated with regulation, cell wall organization, lipid storage, suberization, and other fatty acid metabolic processes (Table **1**).

**Table 1:** List of selected genes highly represented in phellem cells, with predicted role in suberization and/or secondary cell-wall development

| Gene ID | Symbol | Annotation | p-adj | FC |
| --- | --- | --- | --- | --- |
| **Transcriptional regulation** |  |  |  |  |
| AT1G52890 | NAC019 | NAC domain containing protein 19 | 1.76E-08 | 279.98 |
| AT3G24650 | ABI3 | AP2/B3-like transcriptional factor family protein | 1.44E-18 | 105.79 |
| AT2G24430 | NAC038 | NAC domain containing protein 38 | 5.03E-99 | 7.58 |
| AT3G12720 | MYB67 | MYB domain protein 67 | 1.28E-49 | 5.07 |
| AT3G18400 | NAC058 | NAC domain containing protein 58 | 3.33E-76 | 4.46 |
| AT2G36270 | ABI5 | Basic-leucine zipper (bZIP) transcription factor family protein | 1.50E-03 | 2.32 |
| **Lipid storage and lipid localization** |  |  |  |  |
| AT1G05510 | AT1G05510 | naphthalene 1,2-dioxygenase subunit alpha (DUF1264) | 9.27E-10 | 165.89 |
| AT4G25140 | OLEO1 | oleosin 1 | 1.41E-11 | 102.84 |
| AT5G55410 | AT5G55410 | Bifunctional inhibitor/lipid-transfer protein/seed storage 2S albumin superfamily protein | 2.85E-08 | 30.31 |
| AT5G40420 | OLEO2 | oleosin 2 | 4.37E-08 | 26.89 |
| AT2G33380 | RD20 | Caleosin-related family protein | 2.10E-06 | 15.31 |
| AT2G25890 | AT2G25890 | Oleosin family protein | 5.19E-05 | 14.63 |
| AT1G62510 | AT1G62510 | Bifunctional inhibitor/lipid-transfer protein/seed storage 2S albumin superfamily protein | 4.68E-21 | 11.85 |
| AT2G18370 | AT2G18370 | Bifunctional inhibitor/lipid-transfer protein/seed storage 2S albumin superfamily protein | 5.56E-147 | 11.42 |
| AT3G47790 | ABCA8 | ABC2 homolog 7 | 2.51E-40 | 10.16 |
| AT3G58550 | AT3G58550 | Bifunctional inhibitor/lipid-transfer protein/seed storage 2S albumin superfamily protein | 5.18E-116 | 5.63 |
| **Cell wall organization or biogenesis** |  |  |  |  |
| AT5G19890 | AT5G19890 | Peroxidase superfamily protein | 2.11E-05 | 132.76 |
| AT1G65680 | EXPB2 | Expansin B2 | 0.00E+00 | 88.88 |
| AT1G48130 | PER1 | 1-cysteine peroxiredoxin 1 | 1.627E-16 | 44.97 |
| AT5G05340 | PRX52 | Peroxidase superfamily protein | 1.622E-75 | 14.17 |
| AT2G45220 | AT2G45220 | Plant invertase/pectin Methylesterase inhibitor superfamily | 8.13E-140 | 12.87 |
| AT2G18360 | AT2G18360 | alpha/beta-Hydrolases superfamily protein | 1.95E-145 | 12.51 |
| AT5G51500 | AT5G51500 | Plant invertase/pectin methylesterase inhibitor superfamily | 6.97E-03 | 4.58 |
| AT2G18980 | AT2G18980 | Peroxidase superfamily protein | 1.45E-53 | 4.47 |
| AT4G17483 | AT4G17483 | alpha/beta-Hydrolases superfamily protein | 2.40E-11 | 4.35 |
| AT5G37690 | AT5G37690 | SGNH hydrolase-type esterase superfamily protein | 3.27E-65 | 4.30 |
| **Related to suberin biosynthesis** |  |  |  |  |
| AT2G37360 | ABCG2 | ABC-2 type transporter family protein | 1.61E-18 | 8.77 |
| AT3G44550 | FAR5 | fatty acid reductase 5 | 1.75E-136 | 8.10 |
| AT5G63560 | FACT | HXXXD-type acyl-transferase family protein | 1.32E-170 | 7.91 |
| AT3G44540 | FAR4 | fatty acid reductase 4 | 1.01E-129 | 7.57 |
| AT5G58860 | CYP86A1 | cytochrome P450, family 86, subfamily A, polypeptide 1 | 8.36E-68 | 5.32 |
| AT5G13580 | ABCG6 | ABC-2 type transporter family protein | 5.78E-51 | 4.64 |
| AT3G11430 | GPAT5 | glycerol-3-phosphate acyltransferase 5 | 7.17E-41 | 4.32 |
| AT1G53270 | ABCG10 | ABC-2 type transporter family protein | 2.66E-23 | 4.13 |
| **Fatty acid metabolic process** |  |  |  |  |
| AT1G24430 | AT1G24430 | HXXXD-type acyl-transferase family protein | 1.40E-200 | 13.72 |
| AT4G00400 | GPAT8 | glycerol-3-phosphate acyltransferase 8 | 6.36E-96 | 8.09 |
| AT5G08250 | AT5G08250 | Cytochrome P450 superfamily protein | 6.13E-08 | 7.00 |
| AT4G15393 | CYP702A5 | cytochrome P450, family 702, subfamily A, polypeptide 5 | 1.07E-23 | 6.66 |
| AT2G38110 | GPAT6 | glycerol-3-phosphate acyltransferase 6 | 2.63E-51 | 6.08 |
| AT5G42590 | CYP71A16 | cytochrome P450, family 71, subfamily A, polypeptide 16 | 2.42E-04 | 5.55 |
| AT4G15390 | AT4G15390 | HXXXD-type acyl-transferase family protein | 2.81E-55 | 5.23 |
| AT1G49430 | LACS2 | long-chain acyl-CoA synthetase 2 | 1.63E-55 | 4.30 |
| AT4G15400 | BIA1 | HXXXD-type acyl-transferase family protein | 1.08E-05 | 4.17 |
| AT2G45970 | CYP86A8 | cytochrome P450, family 86, subfamily A, polypeptide 8 | 2.79E-34 | 4.14 |
| AT1G78990 | AT1G78990 | HXXXD-type acyl-transferase family protein | 5.78E-29 | 4.05 |

Aiming to highlight the most represented molecular pathways involved in phellem development, we selected the DEGs with a FC > 4. With this threshold, we obtained 977 DEGs in phellem cells (Table **S2**), which were organized in a network, according to predicted protein-protein interactions available on STRING database (Szklarczyk *et al.,* 2019). The generated network retrieved 2,764 interactions across 326 DEGs, and clearly organized in three interconnected clusters (Fig. **4**; Table **S4A**). Genes showing the highest enrichment in phellem cells were mostly found in cluster 2, whereas clusters 1 and 3 comprised genes with lower FC (Fig. **4**). Cluster 1 included genes related to cell wall modifications (Fig. **S5a,** Table **S4B**), namely biosynthesis of diverse cell wall component precursors *(CCoAOMT7, PAL4, 4CL5)*, as well as genes described as specifically involved in suberization pathway *(FAR4/5, CYP86A1, FACT, GPAT5, ABCG2/6)* (Table **1**). Other non-characterized genes involved in lipid metabolism were also identified in this cluster, including HXXXD-type acyl-transferase family genes *(AT4G15390* and *AT1G24430)* and Cytochrome P450s *(AT5G08250, CYP702A5, and CYP71A16)*. Genes associated with lipid transport *(RD20, ABCA8, AT2G18370* and AT3G58550) and diverse genes related to extracellular polymerization were also represented within cluster 1, including α/β-fold protein hydrolases *(AT2G18360 and AT4G17483)*, peroxidases *(AT5G14130, AT5G66390, AT3G32980, AT2G18980, AT2G38390* and *AT2G35380)* and a GDSL esterase *(AT5G37690)* (Table **1**).

**Fig. 4.**
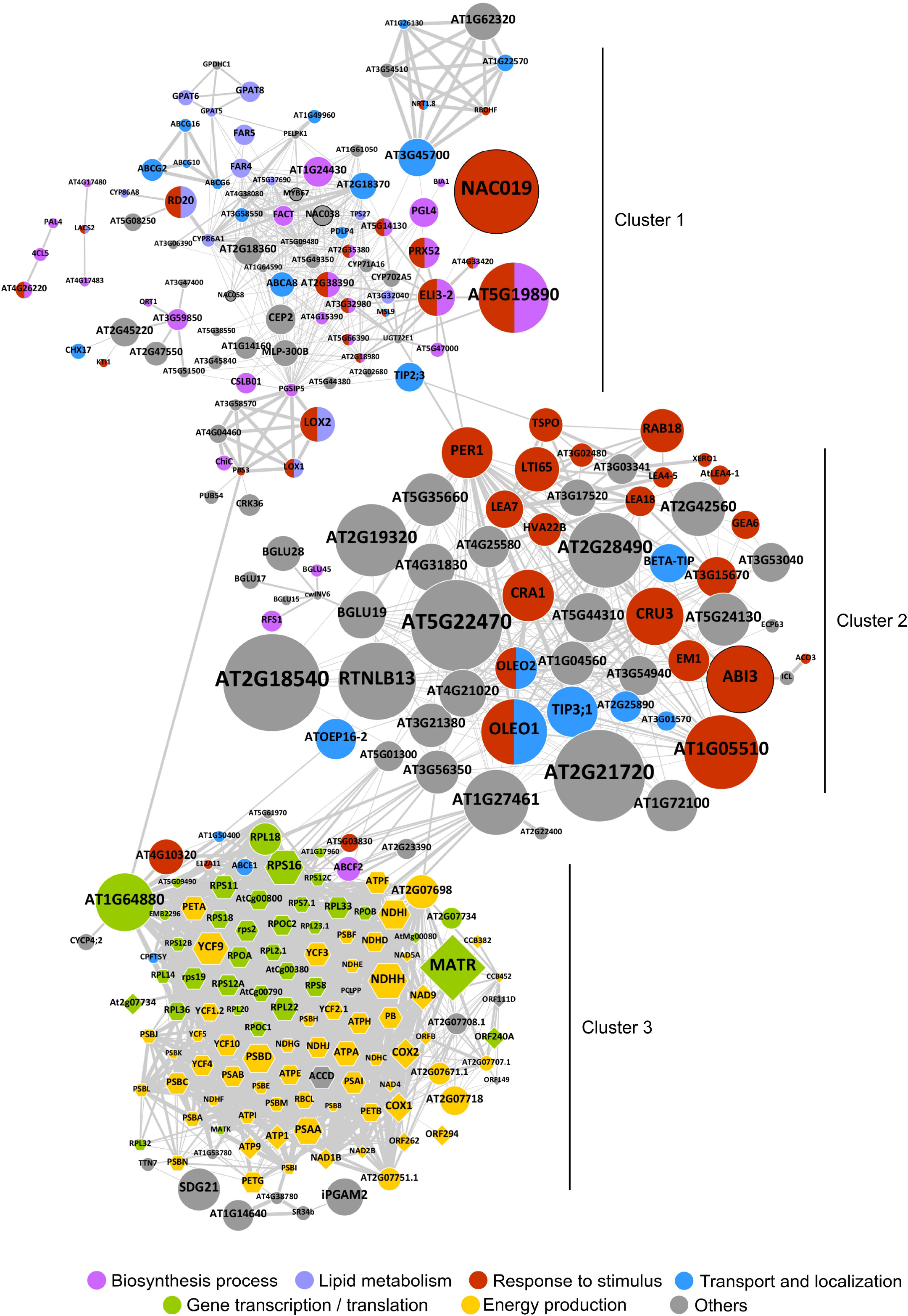
Network representation of predicted protein-protein interactions for DEGs enriched in phellem with FC>4. The network was build based on STRING database with each node representing the protein product associated with each DEG, and each connecting edge representing a predicted association (based on co-expression and inferred or validated protein-protein interactions). Node size corresponds to the Log2FC of each gene and edge thickness represents the confidence value for each association. Circular nodes represent genes from the nuclear genome, while hexagon and diamond shapes represent gene from plastid and mitochondrion genomes, respectively. Fill colors represent general functions or processes annotated to each gene product. Nodes with black border represent TFs.

In cluster 2, most genes were associated with responses to stimulus, including endogenous and external stimuli, and stress (Fig. **S5b,** Table **S4C**). More specifically, DEGs encoding LEA-family proteins were the most abundant group related to response to abiotic stress stimuli with 13 genes identified. In addition, 11 abscisic acid (ABA)-responsive DEGs in phellem were present in this cluster. Interestingly, DEGs coding for proteins related to transport and localization are also grouped in cluster 2. These included water transport *(TIP3;1, BETA-TIP)* and transmembrane transport (*OLEO1*, *OLEO2, ATOEP16-2, AT3G01570, AT2G25890)* (Table **1**).

Finally, in cluster 3 we identified genes mostly related to energy production and transcription or translation activity (Fig. **S5c,** Table **S4D**). Remarkably, this cluster is mainly composed of gene products transcribed from mitochondria and plastid genomes. The grouped DEGs related to energy processes included several elements involved in ATP production, such as ATPases and ATP synthase subunits, NAD(P)H oxidoreductases, electron carriers (cytochrome, ubiquinone and plastoquinone components), and photosystem-related proteins. Besides energy metabolism, a great component of this cluster included DEGs encoding ribosomal subunits (RPL and RPS-related) or related to tRNA activity. DEGs associated with transcription included several plastid-specific RNA polymerases (RPOA, RPOB, RPOC1,-2) and intron maturases (MATK and MATR). In addition, some proteins related to transmembrane transport were also found, such as ABC and TIC transporter proteins and a signal recognition receptor protein *(AT5G61970)*.

### Transcription Factors enriched in phellem

Among the DEGs enriched in phellem (FC > 2), 158 were annotated as TFs, including several families (Table **1**, **S5**). Interestingly, 58 of these TFs are described as responsive to stimulus, of which 23 are involved in ABA signalling or response, including members of diverse TF families as well as central ABA regulators, such as *ABI3* and *ABI5* (Kim, 2014).

Specific TF families are already reported for their involvement in regulatory pathways related to cell wall modifications, in particular WOX, MYB, and NAC families. To explore intrafamily gene diversity and its functional association with phellem development, we performed a phylogenetic analysis of all DEGs from these families identified in our analysis (Fig. **5**). *WOX* TFs are of particular interest, not only because of their predicted role as meristematic regulators, but also because they are specifically expressed during secondary development in both vascular cambium and phellogen (Lehmann and Hardtke, 2016; Jouannet *et al.,* 2015; Furuta *et al.,* 2014; Motte *et al.,* 2019; Smetana *et al.,* 2019). Four *WOX* family members were detected in our analysis, with contrasting expression patterns. While *WOX12* and *WOX2* were enriched in phellem, the closely related *WOX4* and *WOX13* were represented in the *UBQ10*-specific translatome (Fig. **5a**). Phylogenetic analysis of *NAC* and *MYB* TFs expressed in *GPAT5*-specific and *UBQ10*-specific showed association between specific clades and expression pattern. This is the case for two NAC-family clades including the highly enriched *NAC019*, as well as *NAC058* and *NAC38*, which were also previously clustered in STRING analysis together with cell wall modification-related genes (Fig. **5b**, **S5a**). Interestingly, *NTM1, NAC001*, *NAC003* and *NAC048*, which have not been associated with other phellem tissues in previous studies, clustered together in the same clade. Similar associations between clades from MYB-family members and expression pattern were also found (Fig.**5c**). One example is the clade including *MYB9, MYB39, MYB92, MYB93* and *MYB107*, which were previously validated for their role in suberization (Gou *et al.,* 2017; Lashbrooke *et al.,* 2016; Cohen *et al.,* 2020).

**Fig. 5.**
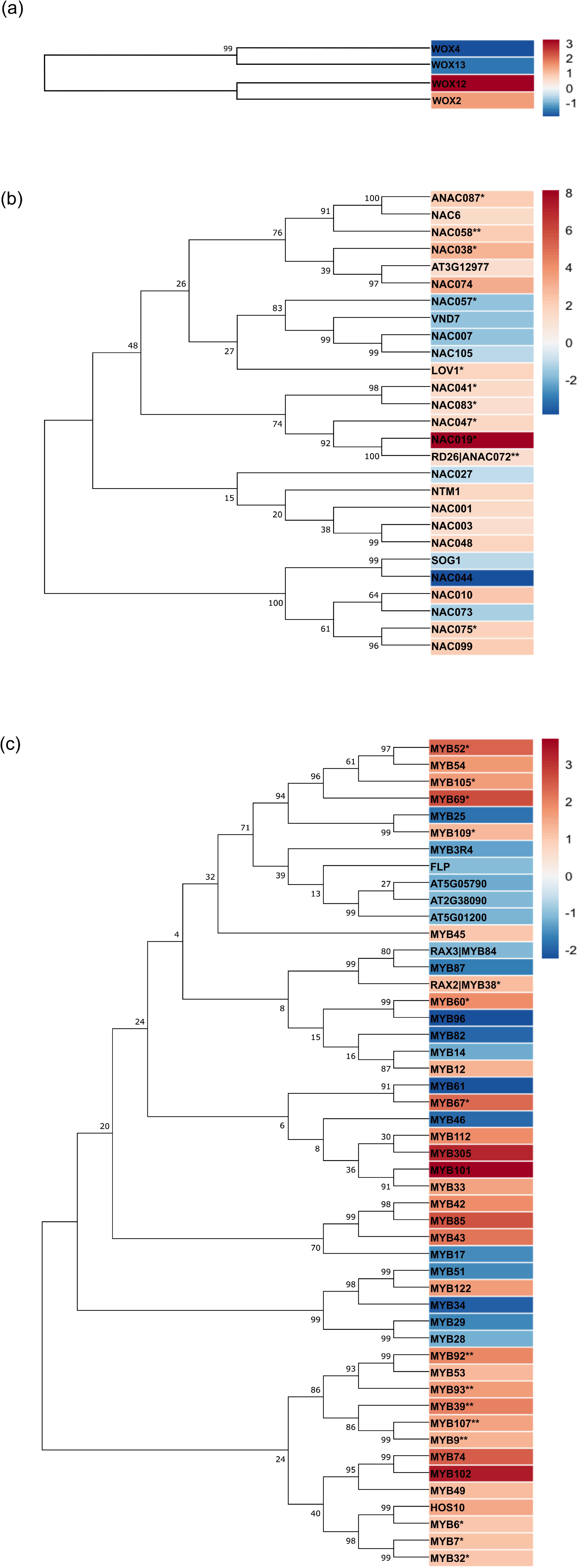
Maximum likelihood phylogenetic trees inferred for WOX (a), NAC (b) and MYB (c) transcription factors identified in this study. Bootstrap values for 500 replications are shown next to the branches. Analyses were conducted in MEGA X. Heatmaps represent the LOG2FC obtained for each gene from the comparison between GPAT5-specific and UBB10-specific translatomes. * phellem enriched TFs also previously identified in other transcriptomic studies (Soler *et al.,* 2007, 2011; Ginzberg *et al.,* 2009; Teixeira *et al.,* 2014, 2018; Vulavala *et al.,* 2019, 2017; Rains *et al.,* 2018; Boher *et al.,* 2018; Alonso-Serra *et al.,* 2019; Lopes *et al.,* 2019); *\*\* TFs with demonstrated function in suberization* (Lashbrooke *et al.,* 2016; Verdaguer *et al.,* 2016; Gou *et al.,* 2017; Soler *et al.,* 2020; Cohen *et al.,* 2020).

## Discussion

### Suberization is an early process in phellem development

We studied the initial steps of phellem differentiation in *Arabidopsis* roots, performing a detailed tracing of the suberization process in newly formed cells during periderm development. Previous transcriptomic studies on cork oak and potato using microdissected periderm samples (including phellogen and early-developed phellem) detected suberization-related genes, suggesting that suberization occurs at an early stage of periderm development. In *Arabidopsis*, the combined monitoring of suberin marker genes activation and anatomical analysis, confirmed that suberization is active right after initiation of periderm development. Activation of suberin biosynthesis is initiated before the rupture and peel-off of epidermis and cortex, at 7 DAG (Fig. **1**). Gene expression is specifically detected after division of the pericycle-daughter cells that dedifferentiate to generate the phellogen. Later on, suberin is detected in these new cells that are gradually becoming the external layer of the root. Altogether, these observations confirm that suberization is initiated early after phellogen establishment and division in the development of *Arabidopsis* root periderm. This corresponds to stages 2 to 3 of root periderm development, as previously defined by Wunderling *et al*. (2018) for the *Arabidopsis* hypocotyl.

### Cell wall modifications characterize the phellem differentiation process

Given its early activation during periderm development, *GPAT5* is here identified as an appropriate marker to track down newly formed phellem cells, and its use for TRAP-SEQ enabled the selection of transcripts enriched in phellem tissue (Fig. **2**). The network analysis of the phellem translatome suggested its organization in at least three main and interconnected regulatory modules. In these modules we found several metabolic processes with a relevant role in phellem development, including synthesis and transport of cell wall components, namely suberin, response to stress and energy production and translation taking place in the plastids.

The most distinctive feature of periderm layers present in different organs and plant-species is suberin deposition in cell walls. In the *Arabidopsis* root phellem we confirmed the expression of multiple genes previously characterized as specific for synthesis of suberin components (e.g. *CYP86A1, CYP86B1, FAR1/4/5, GPAT5/7, FACT, ASFT)*, as well as several genes with predicted functions for a potential role in suberin biosynthesis. These genes include several non-described members of Cytochrome P450 and HXXXD-motif/BAHD acyltransferase families (Table **1**). Moreover, network analysis predicting protein-protein interactions clustered these genes together with suberin biosynthetic genes (Fig. **S5a**) supporting the hypothesis of their involvement in phellem development. In addition, several genes required for the production of suberin precursors were detected as well, which included genes from the phenylpropanoid pathway (*4CL5*, *PAL4* and *AT4G26220*), acyl elongation complexes *(KSC1/2/8, KCR1* and *HCD1)* and various long chain acyl-CoA synthases *(LACS1/2/9/6)*, indicating the capacity of phellem cells in producing precursors needed for suberization (Andersen *et al.,* 2020). In fact, functional enrichment of phellem DEGs highlighted not only suberin biosynthesis but also specific terms related to lipid, fatty acid and phenylpropanoid metabolic processes (Fig. **3**; Table **S3A**).

DEGs related to transport were also particularly enriched in the phellem translatome, including not only the previously characterized suberin ABC transporter members *(ABCG2/6/10/20)* (Yadav *et al.,* 2014), but surprisingly also other strongly represented genes associated with lipid storage and trafficking. Lipid transfer proteins (LTPs), oleophilic bodies and vesicle mediated transport have been proposed to take a part in transport of suberin moieties to the apoplast (Philippe *et al.,* 2020; Hoffmann-Benning, 1994), however, these mechanisms have not been experimentally demonstrated. Our targeted analysis detected for the first-time genes associated with these processes in suberizing tissues, namely several LTPs, oleosins and a caleosin, are enriched in phellem cells (Table **1**). This evidence agrees with the hypothesis that suberin monomers are transported and accumulated in coated oil bodies and LTPs.

Several genes associated with cell wall organization and biogenesis were overrepresented in *Arabidopsis* phellem cells, supporting the described cell wall alterations occurring during phellem differentiation. We detected that specific peroxidases were highly enriched in *Arabidopsis* phellem, and some grouped together in the cell wall modification cluster defined from network analysis (Fig. **S5a**). Peroxidases are described in lignin polymerization (Francoz *et al.,* 2015; Rojas-murcia *et al.,* 2020) and it was hypothesized that such proteins could be also involved in the cross-linking of suberin phenolic monomers to the cell wall components (Bernards *et al.,* 1999; Bernards *et al.,* 2004). In fact several peroxidases are commonly found in the phellem of other plant species (Alonso-Serra *et al.,* 2019; Lopes *et al.,* 2019; Vulavala *et al.,* 2019; Rains *et al.,* 2018; Boher *et al.,* 2018). In addition, GDSL esterase/lipases were identified in phellem, which were recently reported to be involved specifically in suberin biosynthesis and induced by ABA *(AT2G23540, AT5G37690, AT1G74460)* (Ursache *et al.,* 2020), as well as other members which have not been associated with phellem or suberization *(AT1G73610, GLIP6, AT1G7125)*. In parallel, α,β-fold hydrolase are also likely associated with suberin polymerization, and diverse members of this family are identified in our analysis (Table **1**, **S2**), of which *AT1G11090* is for the first-time associated with phellem tissues.

It is likely that the targeted phellem cells were at a stage when major alterations in cell walls were occurring, since genes related to the synthesis of suberin components were found less represented (with lower FC), as compared to genes associated with transport or monomer polymerization and cell wall organization. Synthesis and modification of suberin monomers mostly occur in plastids, corroborating the numerous identified genes associated with protein synthesis in plastids. Moreover, the production of the diverse suberin components requires energy consumption and oxidation power (Bernards, 2002) validating the importance of plastids and mitochondria in the fast process of phellem differentiation.

### Molecular Regulators of Phellem Development

Gene ontology analysis revealed that hormonal control of phellem development mostly involves ABA. This hormone has been described to participate in suberization processes of diverse tissues, by controlling the transcription of diverse suberin biosynthesis enzymes, in control or stress conditions (Boher *et al.,* 2013; Bjelica *et al.,* 2016; Lee *et al.,* 2009; Yadav *et al.,* 2014; Barberon *et al.,* 2016). A possible relation of ABA and phellem formation was previously discriminated in phellem of potato and cork oak, where it was mostly associated with stress-responsive genes (Soler *et al.,* 2007; Vulavala *et al.,* 2019). In fact, the ABA-responsive genes enriched in *Arabidopsis* phellem cells are mostly described in responses to biotic or abiotic stimuli (Fig. **S5b**), which can be a result of the increasing exposure of newly formed phellem to the environment. The detection of *ABI3* and *ABI5* TFs, that are described to function as core regulators in ABA signalling in diverse biological processes (Mary *et al.,* 2003; Skubacz *et al.,* 2016), supports a triggering role of ABA in phellem differentiation. Although with low fold-change, compared with *UBQ10*-specific translatome, several salicylic acid (SA)-responsive genes were found in our analysis (Table **S2**, **S3A**), mostly associated with stress responses. Although the direct involvement in phellem differentiation is not yet established, these genes could be necessary to minimize the effects of oxidative stress, likely to occur during suberin synthesis (Bernards *et al.,* 2004). On the other hand, jasmonic acid (JA) is likely to participate in some aspects of phellem differentiation. Actually, JA-related transcripts have been associated with phellem in different species, and particularly in potato, JA was described to play a central role in healing through periderm development, although with no impact on suberization (Boher *et al.,* 2013; Lulai and Suttle, 2004; Lulai and Suttle, 2009; Lulai *et al.,* 2011). In our analysis, we identify several transcripts associated to JA synthesis and signalling, including *LOX*, *CYP94B3* and *JAZ* (Table **S2**, **S3A**), corroborating an effective activation of JA signalling. The cross-talk of JA with other hormones has been considered crucial in plant responses to stimuli and in developmental processes (Huang *et al.,* 2017; Yang *et al.,* 2019), and it might be that JA acts as a core signal during phellem development, controlling processes from initial differentiation to final cell death.

Several lines of evidence have indicated that although the structure of the diverse plant meristems differs, there are commonalities in the molecular mechanisms underlying their function. Comparisons between lateral meristems and/or apical meristems allowed to identify not only many common regulatory components, such as TFs, cyclins, and receptor-like-kinases, but also specific characteristics for each of the meristems (Schrader *et al.,* 2004; Du *et al.,* 2010; Baucher *et al.,* 2007; Fischer *et al.,* 2019). TFs are important regulatory components in developmental processes, and some TF families have been more associated with cell fate specification. The WOX family was characterized as important in diverse processes of tissue differentiation (Gaillochet and Lohmann, 2015). Some WOX members were indicated as potentially involved in periderm development, namely WOX9 (Lopes *et al.,* 2019; Rains *et al.,* 2018), WOX1 (Rains *et al.,* 2018) and WOX4 (Campilho *et al.,* 2020; Boher *et al.,* 2018; Alonso-Serra *et al.,* 2019), as they were previously found in periderm tissues. In our analysis *WOX12* and *WOX2* were identified as enriched in the phellem, being for the first time, associated to this tissue. *WOX12* induction was previously reported in young roots, specifically in pericycle xylem pole cells, after external auxin application (Liu *et al.,* 2014). *WOX2* was proposed to modulate auxin and cytokinin pathways during stem cell initiation in shoot apical meristem (Zhang *et al.,* 2017; Breuninger *et al.,* 2008). The described activation of these *WOX* genes in differentiation events, in particular the expression in root pericycle cells is of particular interest. Although three auxin responsive genes were also upregulated in *Arabidopsis* phellem (Table **S2**), genes related to auxin were not significantly represented. Regardless of the signalling pathway mediating *WOX* expression, these TFs are potentially operating as regulators of phellem cell fate. Contrasting with previous reports, we identified *WOX4* specifically enriched in *UBQ10-* specific translatome. Although being detected in periderm tissues in multiple studies (Alonso-Serra *et al.,* 2019; Boher *et al.,* 2018), WOX4 is also described to be active during vascular development (Alonso-Serra *et al.,* 2019; Smetana *et al.,* 2019), which likely explains the enrichment in the total root translatome.

Several suberization-related TFs were also represented in *Arabidopsis* phellem cells, including MYB and NAC families, among other families. From the previously characterized MYBs, we detected enrichment of *MYB9, MYB39, MYB93* and *MYB107* in *Arabidopsis* phellem cells. Interestingly these genes are phylogenetically related to *MYB53* that was reported for the first time in phellem. *MYB53* expression is induced by ABA in roots (Gibbs *et al.,* 2014), making it a candidate for ABA-dependent regulation of suberization during phellem development. The phylogenetic analysis performed using all MYB TFs identified in this study agreed with the genome-wide analysis of *Arabidopsis* MYBs performed by Dubos *et al.* (2010), suggesting some degree of functional divergence between the MYBs identified here in two distinct translatomes.

The suberin related *NAC058 (StNAC103* orthologue) and *NAC072/RD26 (StNAC101* orthologue) previouslly described in potato (Verdaguer *et al.,* 2016; Soler *et al.,* 2020) were also identified here to be enriched in the *Arabidopsis* phellem cells. Although these orthologs were never associated with the suberization in *Arabidopsis*, they associate with ABA responses (Coego *et al.,* 2014). Particularly *NAC072/RD26* has a central role in ABA-mediated abiotic stress responses, being also a regulator of senescence processes (Tran *et al.,* 2004; Fujita *et al.,* 2004; Kamranfar *et al.,* 2018). The strongly expressed *NAC019* was previously detected in birch suberized phellem cells (Alonso-Serra *et al.,* 2019), and in *Arabidopsis* it is described to have the same expression pattern as *NAC072/RD26* under dehydration, ABA application and high-salinity (Tran *et al.,* 2004). Our network analysis associated this TF with genes related to cell wall modifications, making it a strong candidate to be involved in the regulation of cell wall differentiation throughout phellem development.

Altogether, our targeted transcriptomic analysis of *Arabidopsis* phellem cells revealed the molecular processes of phellem development, unveiling a variety of regulatory and metabolic processes mostly associated to suberization, lipid transport as well as response to a variety of stimuli. In this work we not only highlighted the conservation of the regulatory and metabolic pathways taking place in phellem from different species, but also revealed new candidates for phellem differentiation. Further functional characterization of these candidates will be promising and necessary for a comprehensive understanding of the molecular mechanisms driving phellem cell fate.

## Experimental procedures

### Plant growth conditions

For *in vitro* assays, seeds were surface sterilized with 70% (v/v) ethanol for 5 min., 20% (v/v) bleach for 5 min. and rinsed 5x with sterile water. Thereafter, seeds were sown on square plates containing sterile half-strength Murashige and Skoog (1/2 MS) medium [0.5 x MS salts, 1% (w/v) sucrose, 0.5 g/L 2-(*N-*morpholino)ethanesulfonic acid MES, 0.1g/L Myo-Inositol, pH 5.7, and 1% (w/v) agar] and stratified at 4 °C for 3 days in the dark. Seeds were germinated on vertically positioned square plates in a growth chamber at 21 °C under continuous light (100 μmol m^-2^ s^-1^).

### Plant material and constructs

*Arabidopsis* thaliana ecotype Columbia (Col-0) was used as wild type and for the transgenic lines used. The lines *pGPAT5::mCITRINE-SYP122, pFAR4::GUS* and *pNF-YC12::GUS* were previously described (Barberon *et al.,* 2016; Domergue *et al.,* 2010; Siefers *et al.,* 2009).

Constructs used for TRAP were generated by Gateway cloning in the pH7m24GW,3 vector (Karimi *et al.,* 2007). The BLRP-FLAG-GFP-RPL18 entry vector was made by overlapping PCR. attB1-BLRP-FLAG-GFP and the RPL18-attB2 sequences were PCR amplified using the primer pairs B1-BLRP-FLAG-GFP_F/ GFP-RPL18_R (with GFP sequence as template) and GFP-RPL18_F/ B2-RPL18_R *(Arabidopsis* gDNA template) (Table S**1**). The final overlapping PCR was performed using the primer pair B1-BLRP-FLAG-GFP_F/ B2-RPL18_R (with previous fragments as template) (Table S1). The BLRP-FLAG-GFP-RPL18 construct was subsequently cloned into a PDONR221 entry vector using BP clonase II (Invitrogen) according to manufactures description. Consecutive expression vectors *(pGPAT5* (2,148 bp before ATG) and *pUBQ10* (1,989 before ATG)-driven) were made by LR-cloning using the LR clonase II enzyme mix (Invitrogen) into a Fasted selection containing R4R2 in destination vector as previously described (Barberon *et al.,* 2016). The corresponding gene identifiers are: *GPAT5*, AT3G11430; *UBQ10*, AT4G05320; *FAR4*, AT3G44540.

### Light microscopy

Histochemical GUS staining was performed on fresh *pFAR4::GUS and pNF-YC12:: GUS* roots as previously described (Beeckman and Engler, 1994). Roots were fixed in a 4% formaldehyde and 1% glutaraldehyde solution in 0.1 M phosphate buffer (pH 7.2), dehydrated and embedded in Technovit 7100 resin (Heraeus Kulzer, Wehrheim, Germany), properly oriented, as previously described (Beeckman and Viane, 2000). Sections were obtained (5 μm), placed on glass slides and stained with 0.05% (w/v) ruthenium red solution. Stained sections were subsequently mounted in DePeX (Sigma – Aldrich) and photographed using a Leica DM6 B upright light microscope. A minimum of 8 plants where analysed for each marker line.

### Confocal microscopy

Confocal laser scanning microscopy imaging was performed either on a Leica SP5 or a Zeiss LSM 710 microscope. Excitation and detection windows were set as follows: Propidium iodide (PI): Ex: 514nm and Em: 650-700 nm; mCITRINE: Ex: 488 nm and Em: 500-550 nm; Fluorol Yellow (FY) and GFP: Ex: 488 nm and Em: 500-550 nm; Nile Red: Ex: 561 nm and Em: 600-700 nm; Calcofluor white: Ex: 405 nm and Em: 430-470 nm.

For FY staining (Brundrett *et al.,* 1991), whole plants were incubated in a freshly prepared solution of 0.01% (w/v) FY (Santa Cruz Biotechnologies) in lactic acid (80%) at 70 °C for 30 min. The samples were washed 3x (5 min. in water) and counter-stained with aniline blue (0.5% (w/v), in water) at room temperature in darkness for 30 min. After staining, plants were washed 3x (10 min. in water) and mounted on 50% glycerol.

For PI staining, plants were incubated for 5 min. in a fresh solution of 15 mM PI, rinsed twice in water and mounted in 50% glycerol.

To image TRAP-lines, plants were cleared using ClearSee protocol (Kurihara *et al.,* 2015), and GFP was detected in combination with two histochemical stainings: Calcofluor White M2R (Sigma-Aldrich) for general cell wall staining, and Nile Red (Sigma-Aldrich) to stain suberin (Ursache *et al.,* 2018). To circumvent the difficulty of the periderm development staging associated to the gradient of secondary growth along the same root, morphological observations were confined to a region of 0.5 cm bellow the hypocotyl-root junction through a time-course experiments from 6 to 10 DAG. A minimum of 10 plants where analysed for all the lines and/or staining described.

### Translating Ribosome Affinity Purification and RNA extraction

Seedlings grown on 1/2 MS agar plates with 1% (w/v) sucrose were sampled at 8 DAG. For translatome profiling, root fragments from the hypocotyl base until the first emerged lateral root were pooled and frozen in liquid nitrogen for three biological replicates per line. Frozen root segments were grinded and homogenized in ice-cold polysome extraction buffer (200 mM Tris-HCl (pH 9.0), 200 mM KCl, 36 mM MgCl_2_, 25 mM ethylene glycol tetraacetic acid, 1 mM dithiothreitol, 50 μg/ml cycloheximide, 50 μg/ml chloramphenicol, 1% Igepal CA-630, 1% Brig-35, 1% Triton X-100, 1% Tween-20, 2% polyoxyethylene (10) tridecyl ether, 1% sodium deoxycholate). After incubation for 10 min. on ice, homogenates were centrifuged at 16,000 ×g at 4 °C for 10 min. to pellet insoluble cell debris. Affinity purification of tagged RPL18-containing polysomes was carried out using anti-FLAG beads (Sigma-Aldrich) incubated with the supernatant at 4 °C for 3 hours. Gels were subsequently collected by centrifugation and washed 4 times with polysome wash buffer (200 mM Tris-HCl, pH 9.0, 200 mM KCl, 36 mM MgCl_2_, 25 mM ethylene glycol tetraacetic acid, 5 mM dithiothreitol, 50 μg/ml cycloheximide, 50 μg/ml chloramphenicol). Polysomes were eluted by resuspension of the washed gel in polysome wash buffer containing 3×FLAG peptide (Sigma-Aldrich). Total RNA was extracted from the final elution using Direct-zol RNA kit (Zymo research). In column DNAse I treatment was performed according to the manufacturer instructions.

### Library preparation and RNA-sequencing

RNA concentration and purity were determined spectrophotometrically using the Nanodrop ND-1000 (Nanodrop Technologies) and RNA integrity was assessed using a Bioanalyser 2100 (Agilent). Per sample, 1 ng of total RNA was used as input for cDNA synthesis using the SMART-Seq v4 Ultra Low Input RNA protocol (version “091817”) from Takara Bio USA, Inc. One ng of purified cDNA was sheared to 300 bp using the Covaris M220 and libraries were prepared with the NEBNext Ultra DNA Library Prep Kit for Illumina (version 7.0 - 3/18), according to the manufacturer′s protocol, and selecting for 250 bp insert size. cDNA-libraries from each sample were equimolarly pooled and sequenced on Illumina NextSeq 500 instrument (v2.5, High Output, 75 bp, Single Reads) at the VIB Nucleomics core (www.nucleomics.be).

### Gene expression analysis

For quantitative PCR analysis, first strand cDNA synthesis was performed using mRNA with an oligo-dT primer using Transcriptor High Fidelity cDNA Synthesis Kit (Roche), according to the manufacturer′s instructions. The cDNA was used as template for amplification by qPCR using gene-specific primers (listed on Table S1). Ubiquitin-conjugating enzyme E2 (At5g25760) was used as internal control. Real Time qPCR was done in a Lightcycler 480 (Roche), using Lightcycler 480 Master I Mix (Roche). Amplification reactions were performed in triplicate for each cDNA sample. Target transcript abundance was calculated according to Pfaffl, 2001.

### Sequence data processing and differential expression analysis

After RNA-sequencing, the resulting FASTQ files were imported into the Galaxy Workflow Environment (Afgan *et al.,* 2018). Quality control of the single strand reads was assessed before and after trimming using FastQC. Trimming of adapters and low quality bases was performed using Trimmomatic with “TruSeq3 (single-ended, for MiSeq and HiSeq)” setting for the adapter sequences to use, and all other parameters by default. Reads were mapped on the *A. thaliana* CDS reference from Araport11 using Salmon (Patro *et al.,* 2017) with default parameters. Finally, tximport was used to summarize transcript-level estimates. To identify differentially expressed genes (DEGs) between tissues, statistical analysis was carried out using the DESeq2 package (Love *et al.,* 2014) in Rstudio 1.1 (http://www.rstudio.com/) with R version 3.5.2 (R Foundation for Statistical Computing., 2018). Expression values were normalized by the “Relative Log Expression” (RLE) normalization. Gene expression was pre-filtered with a threshold of ≥10 counts in at least three samples, and in at least one of the two tissues. Differential expression analysis was performed between *GPAT5*-specific and *UBQ10*-specific samples and DEGs were filtered based on a fold change (FC) ≥ 2 and an adjusted p-value ≤ 0.01 (p-values were corrected using the Benjamini-Hochberg method). DEGs were annotated using Araport11 annotation. The RNA-sequencing data set was deposited in ArrayExpress (http://www.ebi.ac.uk/arrayexpress/experiments/E-MTAB-8949).

Gene Ontology (GO) functional enrichment was performed using BiNGO plugin for Cytoscape version 3.7.1 (Maere *et al.,* 2005). The significant GO terms related to cellular component, biological processes and molecular function were retrieved by applying hypergeometric distribution and adjusted p-value < 0.05.

Functional associations between DEGs in phellem (FC > 4) were predicted using STRING v.11.0 database (Szklarczyk *et al.,* 2019). Confidence score for interactions was set for medium score (above 0.4), taking into account co-expression data and proteinprotein interaction evidences. The gene network obtained from STRING v.11 was imported into Cytoscape version 3.7.1 for further analysis and display.

### Sequence Alignment and Phylogenetic analysis

The MYB, NAC and WOX protein sequences were aligned using ClustalW. Phylogenetic analysis was performed using Maximum Likelihood inference with a JTT substitution model, with 500 bootstrap replicates. Both alignment and phylogenetic analysis were conducted with MEGA X (Kumar *et al.,* 2018).

## Acknowledgements

This work has been financially supported by “Fundação para a Ciência e a Tecnologia” through R&D Unit UID/Multi/04551/2013 and R&D Unit UIDB/04551/2020 (GREEN-IT Bioresources for Sustainability), as well as PhD fellowship PD/BD/113472/2015 (awarded to ARL), Post-Doc fellowship SFRH /BPD/86742/2012 (awarded to PMB) and the Project PTDC/BIA-FBT/29704/2017 (“SuberInStress -Cork formation and suberin deposition: the role of water and heat stress”), funded by FCT/MCTES, and co-financed by FEDER in the scope of POR Lisboa 2020 (also funding researcher HS).

## Author Contribution

ARL, PMB, BP, TB and MMO designed the research; ARL, NV, BP, HS, and TGA performed the experiments; ARL, PMB, BP, TB and MMO analysed and discussed the data; ARL wrote the paper with the help PMB, TB and MMO.

## Notes

### Competing Interest Statement

The authors have declared no competing interest.

