## Supplemental Figures S1 to S5 and Table S1 for "Phellem translational landscape throughout secondary development in *Arabidopsis* roots"

The following Supporting Information is available for this article:

**Fig. S1** Development and suberization of periderm tissue in *Arabidopsis* thaliana roots.

**Fig. S2** Validation of differential expression levels in phellem (pGPAT5-/pUBQ10-specific translated RNA) for 5 selected genes.

**Fig. S3** Activation of *NF-YC12* gene during periderm development of *Arabidopsis* thaliana roots.

**Fig. S4** Representative GO terms significantly enriched within genes differentially expressed in total root samples.

**Fig. S5** Detailed representation of the different clusters identified from network for predicted associations of phellem-enriched DEGs.

**Table S1** List of primers used in overlapping PCR and in RT-qPCR.

**Table S2** List of differentially expressed genes with adjusted p-value < 0.01 and |FC| > 2 of phellem and total fraction libraries.

**Table S3A** Functional enrichment analysis showing GO terms significantly enriched within genes differentially expressed in phellem fraction (GPAT5-specific).

**Table S3B** Functional enrichment analysis showing GO terms significantly enriched within genes differentially expressed in total fraction (UBQ-10-specific).

**Table S4A** Network analysis of phellem enriched genes based on protein-protein interactions available in STRING database.

**Table S4B** List of phellem enriched genes displayed in Cluster 1, after STRING analysis.

**Table S4C** List of phellem enriched genes displayed in Cluster 2, after STRING analysis.

**Table S4D** List of phellem enriched genes displayed in Cluster 3, after STRING analysis.

**Table S5** List of differentially expressed TF genes and TF families in phellem cells.

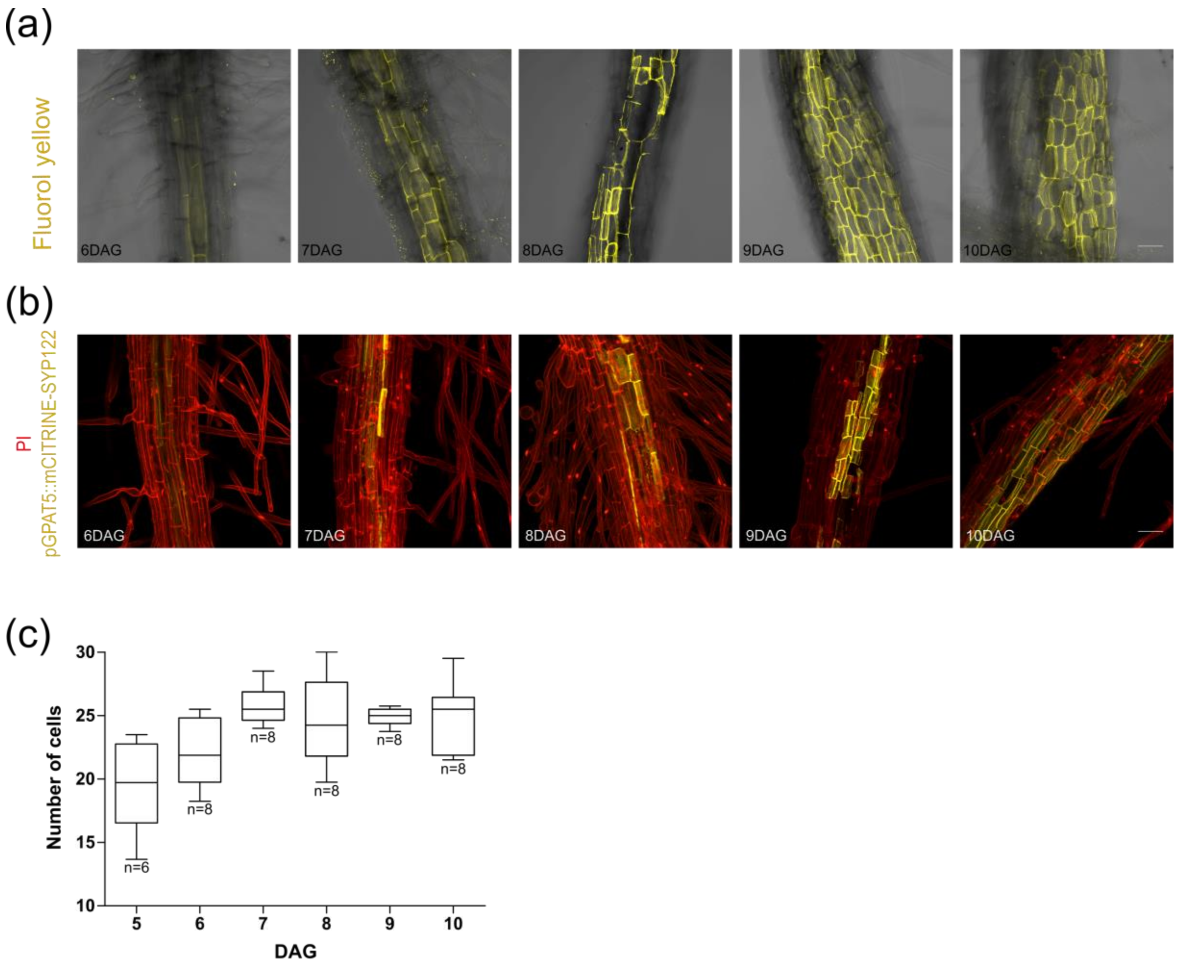


***Fig. S1*** *Development and suberization of periderm tissue in Arabidopsis thaliana roots. (a) Fluorol yellow staining of suberin (yellow) in primary roots. (b) GPAT5::mCITRINE-SYP122 expression (yellow) and propidium iodide staining (red) of primary roots. Images were taken in the periderm development region, below root-hypocotyl junction, between 6 and 10 DAG. The 3D maximum projection figures were obtained by confocal laser scanning with Z-stack images. Scale bar: 50 µm. Images are representative of all analysed samples. (c) Number of circumference cells in the stele (from pericycle to phellem) from 5 to 10 DAG. Cell number was counted in plastic-embedded pFAR4::GUS sections (representatives in Fig. 1a). Middle line of the boxplot is median for n ≥ 6 plants.*

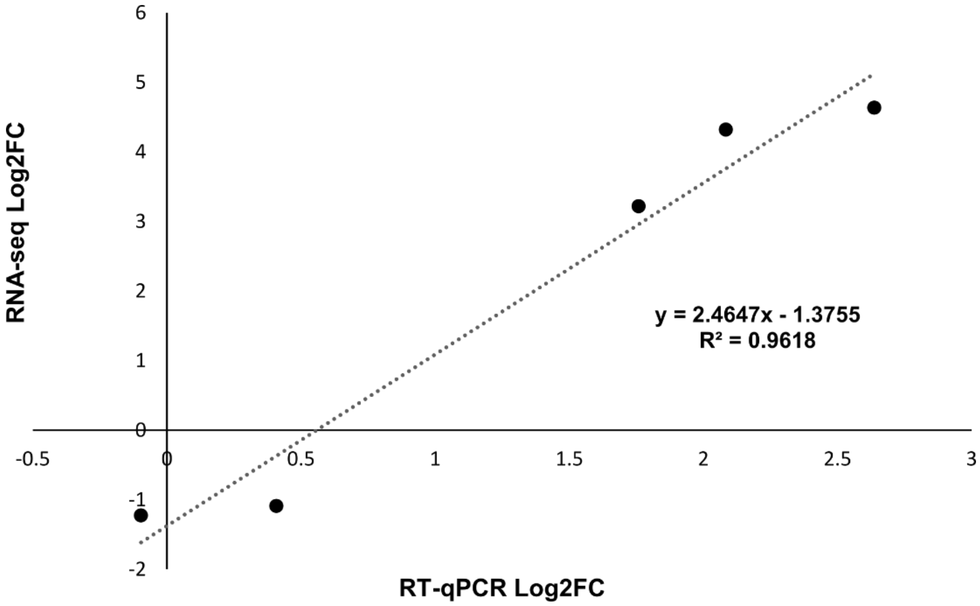


***Fig. S2*** *Validation of differential expression levels in phellem (pGPAT5-specific / pUBQ10-specific RNA) for 5 selected genes. Linear regression with the correlation coefficient ($R^2$). RNA used in RT-qPCR and in RNA-seq was obtained by TRAP.*

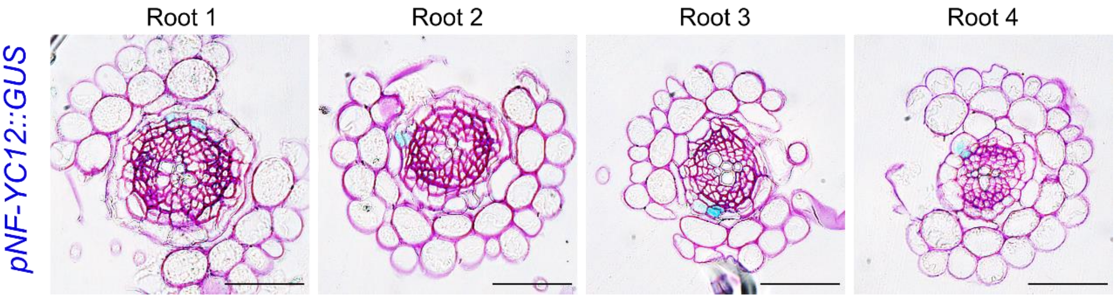


***Fig. S3*** *Activation of NF-YC12 gene during periderm development of Arabidopsis thaliana roots. Plastic embedded cross-sections of pNF-YC12::GUS, taken below the hypocotyl-root junction between 7 and 8 DAG. Four independent representative roots are shown. Scale bar: 50 µm. NF-YC12 was previously detected to be expressed in primary developed root tips and in leaf cells* (Siefers *et al.*, 2009). *In this work, when observations are targeted to the secondary developed root, we detected punctual expression of NF-YC12 in new phellem cells. NF-Y transcription factors (also known as CCAAT box binding factors -CBFs) are discussed to be important regulators of numerous plant developmental and stress-induced responses. NF-YCs are sequence-specific transcription factors with histone-like subunits and are described to form heterotrimeric complexes composed of single subunits from each of three protein families: NF-YA, NF-YB, and NF-YC* (Laloum *et al.*, 2013; Zhao *et al.*, 2017).

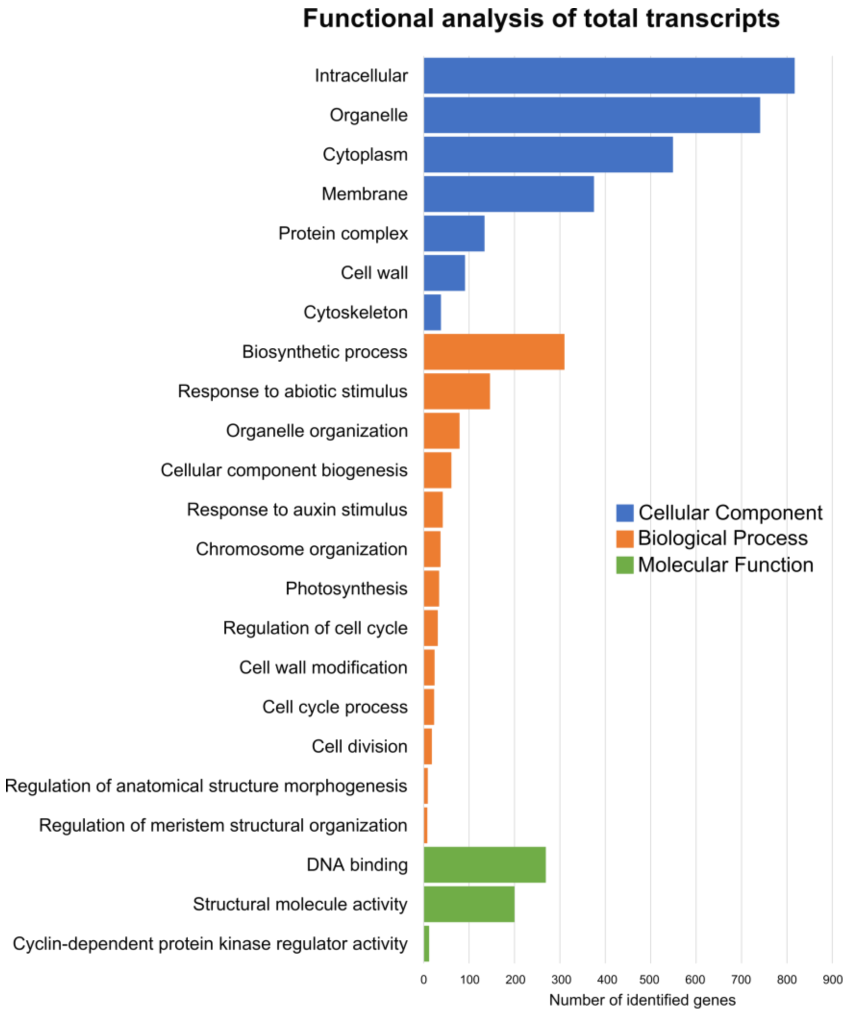


***Fig. S4*** *Representative GO terms significantly enriched within genes differentially expressed in total root samples. Colored bars represent the number of genes in the selected GO terms in Cellular component, Biological process and Molecular function.*

(a)

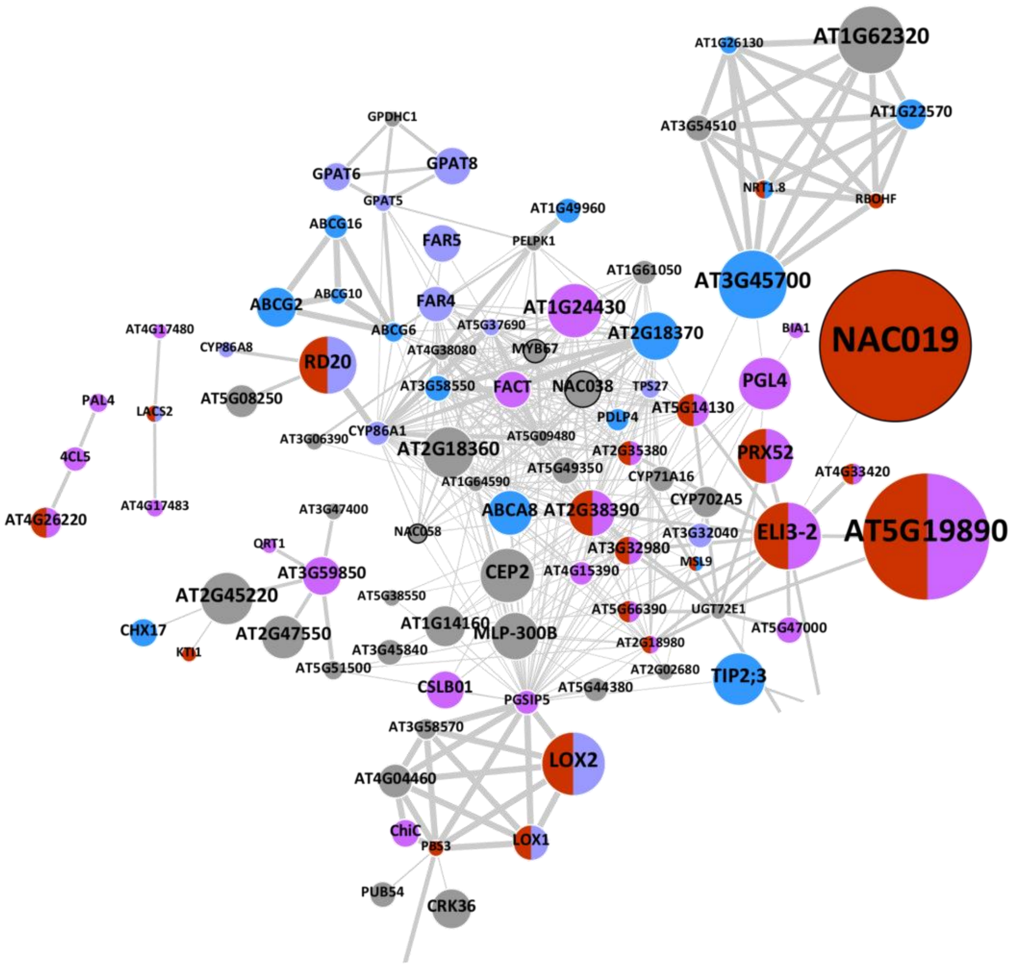

AT1G26130
AT1G62320
AT1G22570
AT3G54510
NRT1.8
RBOHF
GPDHC1
GPAT6
GPAT8
GPAT5
AT1G49960
ABCG16
FAR5
PELPK1
AT1G61050
AT3G45700
ABCG2
ABCG10
FAR4
AT1G24430
AT5G37690
AT2G18370
BIA1
NAC019
AT4G17480
CYP86A8
RD20
ABCG6
AT4G38080
MYB67
AT3G58550
FACT
NAC038
TPS27
PGL4
PAL4
LACS2
AT5G08250
AT5G14130
PDLP4
AT3G06390
CYP86A1
AT2G18360
AT5G09480
AT2G35380
PRX52
4CL5
AT5G49350
CYP71A16
AT4G33420
AT1G64590
AT4G26220
AT4G17483
AT3G47400
ABCA8
AT2G38390
CYP702A5
AT5G19890
NAC058
AT3G32040
ELI3-2
QRT1
AT3G32980
MSL9
AT3G59850
CEP2
AT4G15390
AT2G45220
AT5G38550
AT5G66390
UGT72E1
CHX17
AT1G14160
AT2G47550
MLP-300B
AT5G47000
KTI1
AT3G45840
AT2G18980
AT5G51500
AT2G02680
TIP2;3
CSLB01
AT5G44380
PGSIP5
AT3G58570
LOX2
AT4G04460
ChiC
PBS3
LOX1
PUB54
CRK36

(b)

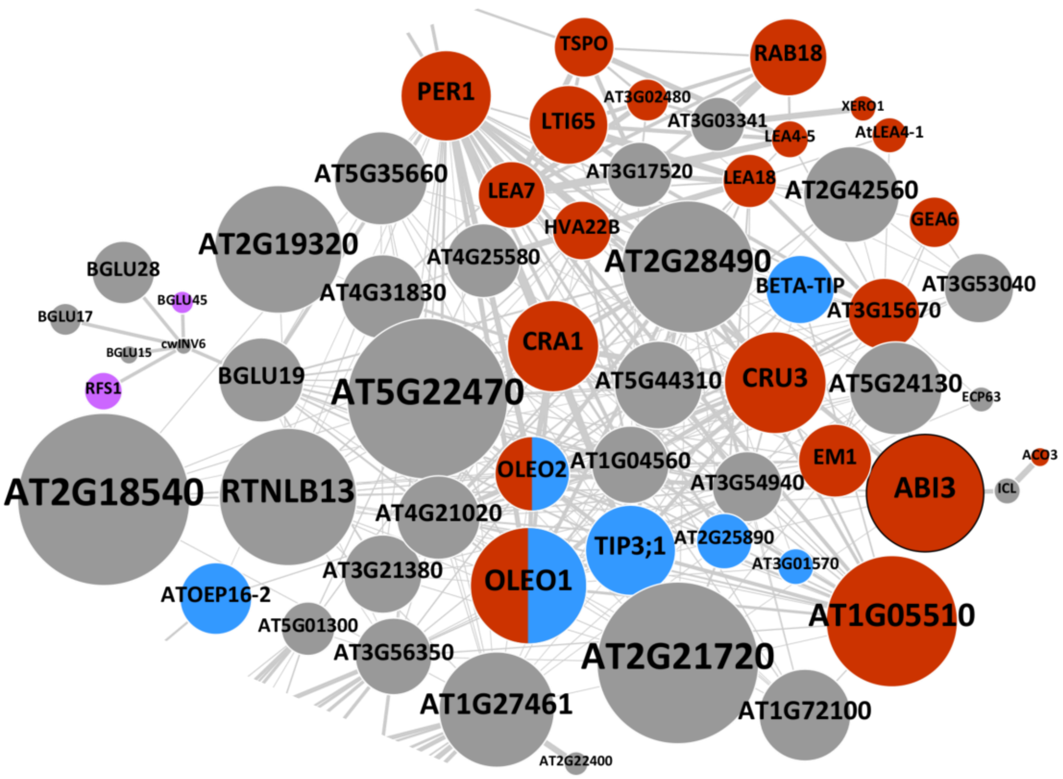

TSPO
RAB18
PER1
AT3G02480
XERO1
LTI65
AT3G03341
LEA4-5
AtLEA4-1
AT5G35660
AT3G17520
LEA18
AT2G42560
LEA7
GEA6
HVA22B
AT2G19320
AT4G25580
AT2G28490
BGLU28
BGLU45
BETA-TIP
AT3G53040
AT4G31830
AT3G15670
BGLU17
cwINV6
BGLU15
CRA1
RFS1
BGLU19
AT5G44310
CRU3
AT5G24130
AT5G22470
ECP63
EM1
ACO3
OLEO2
AT1G04560
AT2G18540
RTNLB13
AT3G54940
ABI3
ICL
AT4G21020
TIP3;1
AT2G25890
AT3G01570
AT3G21380
OLEO1
ATOEP16-2
AT1G05510
AT5G01300
AT3G56350
AT2G21720
AT1G27461
AT1G72100
AT2G22400

(c)

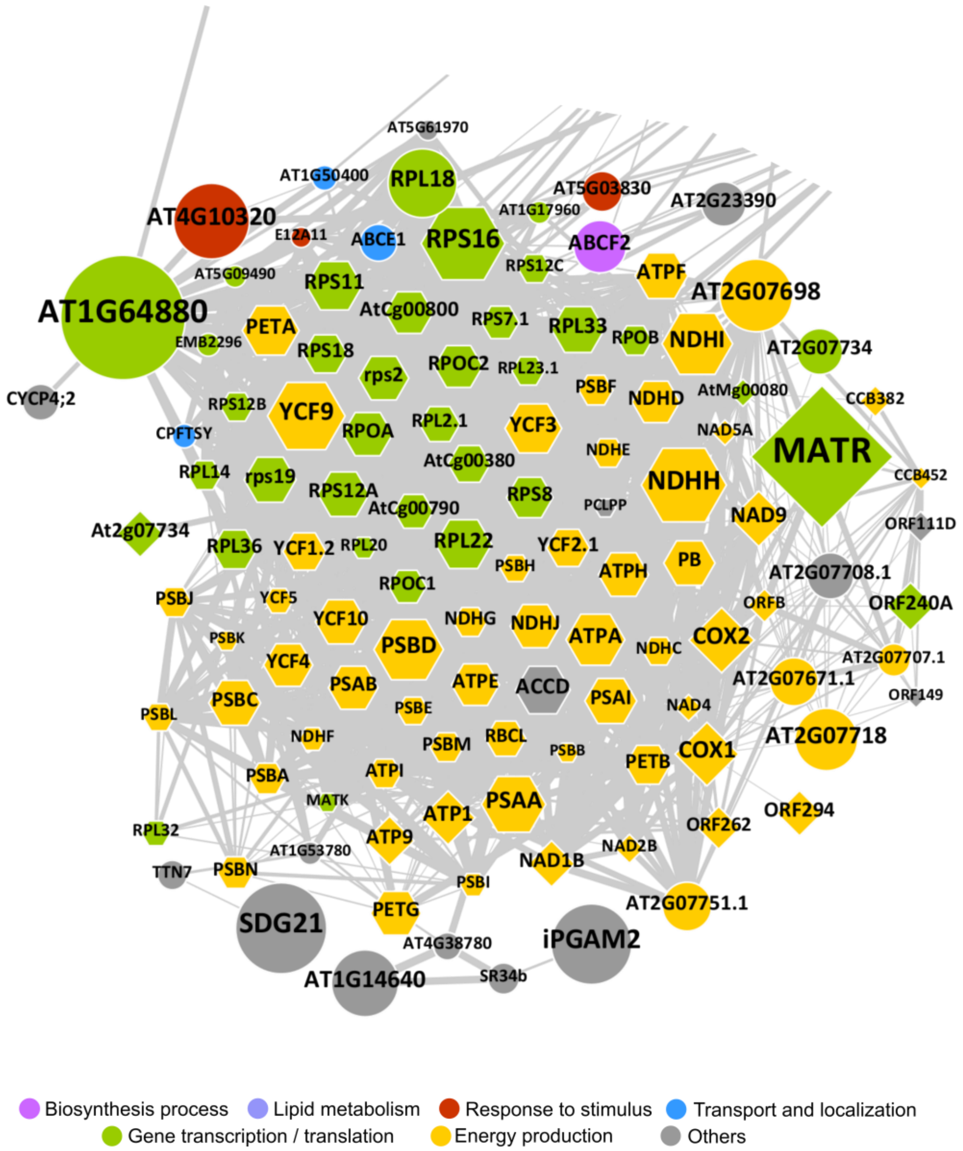


***Fig. S5*** *Detailed representation of the different clusters identified from network for predicted associations of phellem-enriched DEGs. Protein identifiers are attributed to each network node to cluster 1 (a), cluster 2 (b) and cluster 3 (c). Node size corresponds to the Log2FC of each gene and thickening of connecting edges represent the confidence value for the association. Circle nodes represent genes present in the nuclear genome, while hexagon and diamond shapes represent genes present in plastid and mitochondrion genomes, respectively. Fill colors represent functions or processes attributed to each gene product. Nodes with black border represent TFs.*

**Table S1** List of primers used in overlapping PCR and in RT-qPCR.

| Primer name | Sequence (5'-3') | Description |
|---|---|---|
| B1-BLRP-FLAG-GFP_F | AAAAAGCAGGCTTAATGGGACTTAACGATATCTTCGAAGCTCAGAAGATTGAATGGCATGGAGGTGATTATAAGGATGATGATGATAAGGGAGGTGTGAGCAAGGGCGAGG | Primers used for overlapping PCR. BLRP - Biotin Ligase Recognition Peptide; FLAG - DYKDDDDK-tag; GFP-Green Fluorescence Protein; RPL18 – Ribosomal Protein L18. |
| GFP-RPL18_R | GATCAATACCGGACATCTTGTACAGCTCGTCC | |
| GFP-RPL18_F | GGACGAGCTGTACAAGATGTCCGGTATTGATC | |
| B2-RPL18_R | AGAAAGCTGGGTTTAAACCTTGAATCCACGACTCTTCC | |
| AtGPAT5_qPCR1_F | GGGTTTGAGTGCACCAACTT | Primers used for RT-qPCR. The corresponding gene identifiers are: *GPAT5* - AT3G11430; *ASFT*- AT5G41040; *ABCG6* - AT5G13580; *UBCE2* - AT5G25760; *CKA,1* - AT3G48750; *CKA2* - AT3G50000. |
| AtGPAT5_qPCR1_R | GGAGACAAGGCTCGAAAGTG | |
| AtASFT_qPCR1_F | GGTCAAACCTGAATCCGAGA | |
| AtASFT_qPCR1_R | CTTGGACTGCTTCCTCGTTC | |
| AtABCG6_qPCR1_F | ATGAACCAACTTCGGGTCTG | |
| AtABCG6_qPCR1_R | CGGGACAAGAAGAGAAGACG | |
| AtUBCE2_qPCR1_F | CTTGGACGCTTCAGTCTGTG | |
| AtUBCE2_qPCR1_R | GGCGAGGCGTGTATACATTT | |
| CDKA;,1_qPCR1_F | ATTGCGTATTGCCACTCTCATAGG | |
| CDKA,;1_qPCR1_R | TCCTGACAGGGATACCGAATGC | |
| CKA2_qPCR1_F | ACCACCATTAACGTGCGTCAAC | |
| CKA2_qPCR1_R | GATCTTGGCGAGAGAATCGGTATC | |

**Table S2** to **Table S5** are in supplemental Excel files.
